# Marine sulfated glycans inhibit the binding of *Borrelia burgdorferi* adhesins to heparin/GAGs

**DOI:** 10.64898/2026.09.22.753655

**Authors:** Changkai Bu, Ke Xia, Carly Fernandes, Yi-Pin Lin, Nikhat Parveen, Vitor H. Pomin, Jonathan S. Dordick, Lianli Chi, Fuming Zhang

**Author notes:** Correspondence authors. Email addresses (F. Zhang), and (L. Chi).

## Abstract

Recognition of host glycosaminoglycans (GAGs) is critical for the adhesion and colonization of *Borrelia burgdorferi*, but the structural requirements for these interactions remain incompletely understood. Using surface plasmon resonance (SPR), we characterized the heparin-binding properties of three *Borrelia* adhesins, i.e. Bgp, DbpA-A9, and DbpB-B31 and evaluated inhibitors comprising heparin oligosaccharides, selectively desulfated heparins, mammalian GAGs, marine sulfated glycans, pentosan polysulfate (PPS), and mucopolysaccharide polysulfate (MPS). All three adhesins bound directly to heparin, with equilibrium dissociation constants (*K_D_*) of 2.18 nM for Bgp and 36.4 nM for DbpB-B31. Competition assays identified *N-*sulfation and 6-*O*-sulfation as major determinants of binding, whereas 2-*O*-sulfation had a lower effect. Bgp showed chain-length dependence over dp4-dp20, whereas DbpB-B31 preferentially recognized longer oligosaccharides, particularly those of dp12 or greater. Chondroitin sulfate E also exhibited strong inhibitory activity, indicating that recognition was not restricted to heparin/heparan sulfate backbones. Among the tested sulfated glycans, fucosylated chondroitin sulfate (HfFucCS) from *Holothuria floridana* was the most potent inhibitor of Bgp, with an IC_50_ of approximately 4 ng/mL, whereas PPS was the strongest inhibitor of DbpB-B31, followed by MPS. These findings reveal protein-specific sulfated-glycan recognition and identify promising candidates for disrupting *B. burgdorferi*-host GAG interactions.

## 1. Introduction

Lyme disease is one of the most significant tick-borne bacterial infections in temperate regions of the Northern Hemisphere, with an increasing incidence and expanding geographic distribution driven by changes in tick range, climate, land use, wildlife-host ecology, and human exposure. It is caused by members of the *Borrelia burgdorferi* sensu lato complex (Kugeler et al., 2021; Lantos et al., 2021). In the United States alone, approximately 476,000 people are estimated to be diagnosed and treated for Lyme disease annually (Kugeler et al., 2021). Although Lyme disease can generally be treated effectively with antibiotics, particularly when therapy is initiated early, early manifestations may be subtle or nonspecific, delaying diagnosis and treatment. Moreover, currently available interventions rely largely on antibiotic therapy after infection has become established, underscoring the need for new preventive and therapeutic strategies that target the earliest stages of host infection. In particular, agents capable of interfering with spirochetal attachment and tissue colonization could provide a complementary approach to conventional antimicrobial therapy. Transmission of *B. burgdorferi* generally requires prolonged attachment of an infected *Ixodes* tick, with transmission risk increasing with the duration of tick feeding. Following transmission, spirochetes initially encounter host cells and extracellular matrix components at the bite site before disseminating to distal tissues (Eisen, 2018). This early stage of infection may therefore provide an important window for locally administered prophylactic or therapeutic intervention.

During blood feeding, *B. burgdorferi* migrates from the tick midgut to the salivary glands and is subsequently transmitted into the host skin through tick saliva. Tissue dissemination and colonization are facilitated by interactions between bacterial adhesins and host extracellular-matrix components or cell-surface receptors. Its pathogenicity does not primarily depend on toxins or acute tissue destruction. Instead, the spirochete uses its motility, surface lipoproteins, antigenic variation, and diverse adhesins to disseminate through the skin, vascular endothelium, extracellular matrix, facilitate colonization of different organs and connective tissues (Harman et al., 2012; Moriarty et al., 2008; J. R. Zhang, Hardham, Barbour, & Norris, 1997). This process promotes persistent infection and elicits inflammatory and immune responses that contribute to disease manifestations. Among these host ligands, glycosaminoglycans (GAGs), including heparan sulfate (HS), dermatan sulfate (DS), and chondroitin sulfate (CS), serve as structurally diverse, negatively charged binding molecules for *B. burgdorferi*. Several *B. burgdorferi* surface proteins have been shown to mediate attachment to GAGs or GAG-bearing proteoglycans (Leong, Morrissey, Ortega-Barria, Pereira, & Coburn, 1995; Leong, Robbins, Rosenfeld, Lahiri, & Parveen, 1998).

GAGs are linear, negatively charged polysaccharides composed primarily of repeating disaccharide units. They are major components of the cell surface and ECM and regulate numerous molecular interactions in these environments. Heparan sulfate (HS), one of the most abundant GAGs in mammalian tissues, is covalently attached to core proteins to form heparan sulfate proteoglycans (HSPGs), which participate in diverse biological processes, including receptor activation, signal transduction, cytoskeletal remodeling, and intercellular communication (Sarrazin, Lamanna, & Esko, 2011). Because pathogen adhesion to host tissues often depends on specific protein-glycan interactions, soluble decoy molecules that mimic sulfated host glycans may provide a strategy for interfering with these interactions (Rostand & Esko, 1997).

*B. burgdorferi* expresses multiple GAG-binding adhesins, including Bgp, DbpA, and DbpB (Fischer, Parveen, Magoun, & Leong, 2003; Guo, Brown, Dorward, Rosenberg, & Hook, 1998; Parveen & Leong, 2000). Bgp is a multifunctional surface protein that can be shed into the extracellular environment under adverse growth conditions and exhibits nucleosidase activity (Cluss, Silverman, & Stafford, 2004; Parveen et al., 2006). Determining whether individual adhesins preferentially recognize specific sulfated-polysaccharide structures, and how sulfation patterns, chain length, and the spatial distribution of sulfate groups, may provide a structural basis for developing novel anti-adhesion strategies (Leong et al., 1998). Marine-derived sulfated polysaccharides possess diverse glycan backbones, sulfation patterns, and spatial charge distributions that contribute to a broad range of biological activities, including anticoagulant, antiviral, anti-inflammatory, and immunomodulatory effects (Dwivedi et al., 2021; Pomin & Mourao, 2008; Yang, Song, Xia, et al., 2024; F. Zhang et al., 2022). These polysaccharides have therefore attracted increasing interest as GAG mimetics and potential inhibitors of pathogen adhesion (Shi et al., 2023; Song et al., 2021; Yang, Song, Jin, et al., 2024).

In this study, we assembled a glycan library (Fig. 1) comprising heparin and structurally diverse marine sulfated polysaccharides and used surface plasmon resonance (SPR) to systematically evaluate their ability to inhibit the binding of *B. burgdorferi* GAG-binding adhesins to immobilized heparin. Comparison of the inhibition profiles of Bgp and DbpB-B31 revealed distinct preferences for different sulfated polysaccharides. These findings indicate that adhesion-glycan recognition is governed not solely by overall negative charge but also by structural features such as the glycan backbone, sulfation pattern, chain length, and spatial distribution of sulfate groups. Our results identify marine sulfated polysaccharides as potential competitive inhibitors of adhesion-heparin interactions and provide a basis for evaluating sulfated glycans as GAG-based decoys for pathogen attachmen, which may provide a foundation for developing glycan-based anti-adhesion strategies for Lyme disease.

**Fig. 1.**
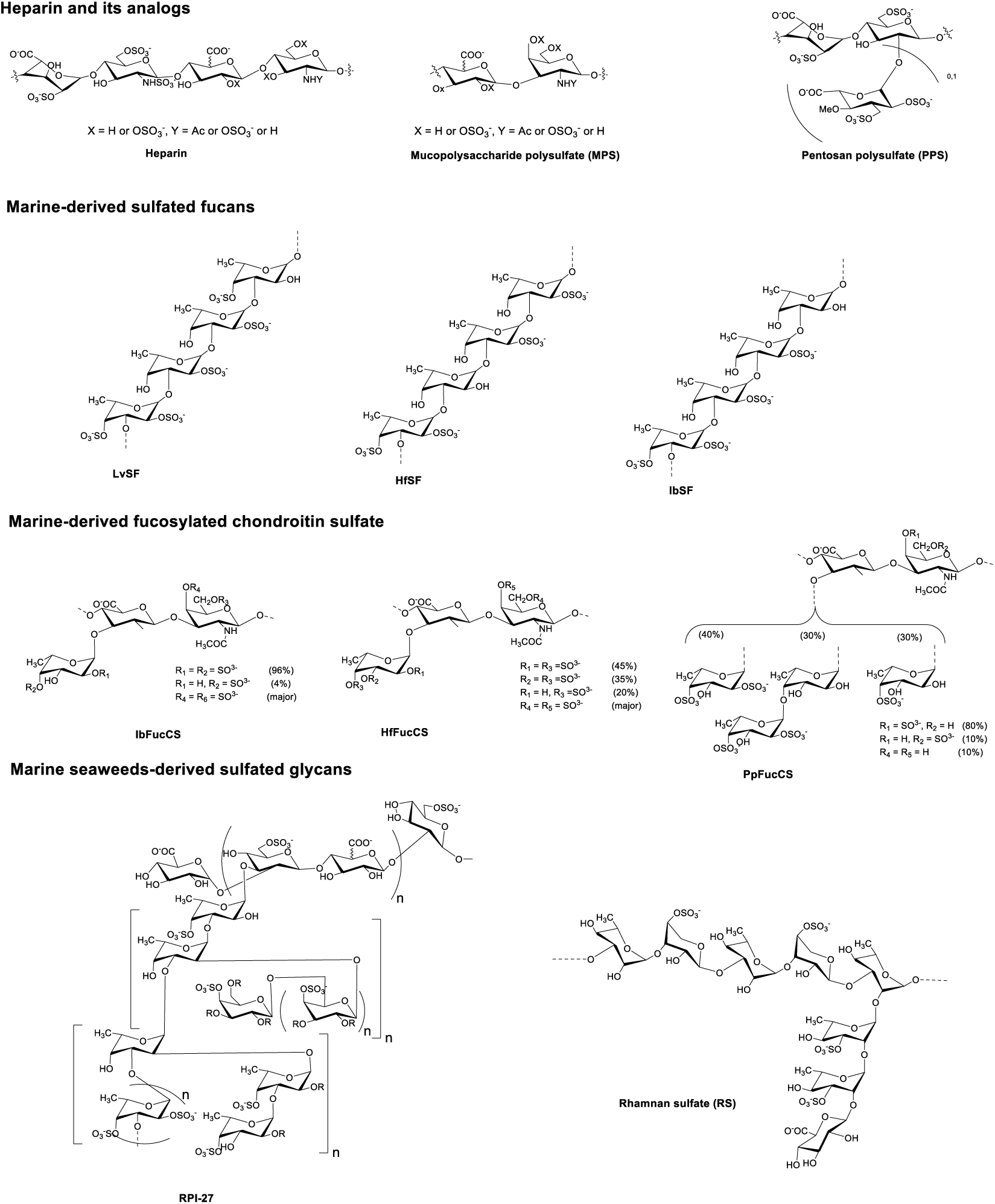
Chemical structures of selected sulfated glycans

## 2. Materials and methods

### 2.1. Materials

Porcine intestinal heparin (average molecular weight, 15 kDa) and heparan sulfate (HS; average molecular weight, 12 kDa) were purchased from Celsus Laboratories (Cincinnati, OH, USA). 6-*O*-Desulfated heparin (6-DeS heparin; MW, 13 kDa) was provided by Dr. Wang from the University of South Florida. 2-*O*-Desulfated IdoA heparin (2-DeS heparin; MW, 13 kDa), *N*-desulfated heparin (*N*-DeS heparin; MW, 14 kDa) and heparin oligosaccharides with varying degrees of polymerization (dp4 to dp20) were purchased from Iduron (Manchester, UK). Chondroitin sulfate A (CSA; 20 kDa) from porcine rib cartilage, dermatan sulfate (DS; 30 kDa) from porcine intestine, and chondroitin sulfate C (CSC; 20 kDa) from shark cartilage were purchased from Sigma-Aldrich (St. Louis, MO, USA). Chondroitin sulfate D (CSD; 20 kDa) from whale cartilage and chondroitin sulfate E (CSE; 20 kDa) from squid cartilage were purchased from Seikagaku (Tokyo, Japan). Keratan sulfate (KS; 14.3 kDa) was isolated from bovine cornea in the Linhardt laboratory.

Marine-derived sulfated glycans, including IbSF (∼100 kDa), IbFucCS (70–80 kDa), PpFucCS (∼60 kDa), LvSF (∼100 kDa), HfSF (∼100 kDa), and HfFucCS (60–100 kDa), were isolated from the sea cucumbers *Isostichopus badionotus*, *Holothuria floridana*, and *Pentacta pygmaea*, as well as the sea urchin *Lytechinus variegatus*, in Dr. Pomin’s laboratory at the University of Mississippi (Dwivedi et al., 2022). Mucopolysaccharide polysulfate (MPS; 14.5 kDa) was obtained from Luitpold Pharma (Munich, Germany), and pentosan polysulfate (PPS; 6.5 kDa) was obtained from Bene Pharma (Munich, Germany). RPI-28 (fucoidan) was purified from the brown seaweed *Saccharina japonica* in Dr. Weihua Jin’s laboratory (Kwon et al., 2020). Rhamnan sulfate (RS) was purified from the green seaweed *Monostroma nitidum* in Zhang’s lab (Song et al., 2021). The chemical structures of the sulfated glycans are shown on Fig.1.

Recombinant Bgp protein was produced using an *E. coli* expression system. The pET30a-Bgp expression plasmid was provided by Dr. Nikhath Parveen. Recombinant DbpA-A9, and DbpB-B31 proteins were provided by Dr. Yi-Pin Lin’s lab. Streptavidin (SA) sensor chips used in this study were purchased from Cytiva (Uppsala, Sweden). SPR measurements were performed using a Biacore T200 SPR instrument (Cytiva, Uppsala, Sweden), and the data were processed using Biacore T200 Evaluation Software (version 3.2).

### 2.2. Heparin chip preparation

Biotinylated heparin was synthesized as follows. First, 1 mg of heparin and 1 mg of amino-PEG3-biotin (Thermo Scientific, Waltham, MA, USA) were dissolved in 200 μL of water. Subsequently, 5 mg of NaCNBH₃ was added to the solution, and the mixture was incubated at 70 °C for 24 h. An additional 5 mg of NaCNBH₃ was then added, and the reaction was continued for another 24 h. Upon completion of the reaction, the mixture was desalted using a centrifugal filter with a molecular weight cut-off of 3 kDa, and the resulting biotinylated heparin was lyophilized for subsequent chip preparation. To prepare heparin chip, biotinylated heparin (0.1 mg/mL in HBS-EP+ buffer: 0.01 M HEPES, pH 7.4, 0.15 M NaCl, 3 mM EDTA, and 0.05% v/v surfactant P20; Cytiva, Uppsala, Sweden) was injected at a flow rate of 10 μL/min over flow cells 2, 3, and 4 of an SA sensor chip. Flow cell 1 was immobilized with biotin only and served as the reference channel.

### 2.3. Study of the binding kinetics and affinity of B. burgdorferi GAG-binding proteins to heparin

Recombinant Bgp, DbpA-A9, and DbpB-B31 proteins for SPR analysis were diluted to a series of concentrations using HBS-EP+ running buffer. Protein samples were injected over the sensor surface at 25 °C at a flow rate of 45 μL/min. The association phase lasted 120 s, followed by a 180 s dissociation phase using running buffer. After dissociation, 45 μL of 2 M NaCl was injected over the chip surface for regeneration.

### 2.4. SPR analysis of competitive inhibition by heparin oligosaccharides and chemically modified heparins

Solution competition SPR experiments were used to compare the effects of heparin oligosaccharides and selectively desulfated heparins, on the binding of Bgp and DbpB-B31 to immobilized heparin. Bgp and DbpB-B31, at concentrations of 37.5 nM and 675 nM, respectively, were pre-mixed with 10 µM heparin, heparin oligosaccharides of varying degrees of polymerization (dp4-dp20), or selectively desulfated heparin. The mixtures were injected over a heparin-coated chip at 25 °C at a flow rate of 45 μL/min. The chip surface was regenerated after each cycle by injecting 45 μL of 2 M NaCl. Samples containing protein alone, without soluble competitors, served as non-competition controls. Their response values were defined as 100% and used to calculate the relative binding response for each competitor.

### 2.5. SPR analysis of competitive inhibition by different GAGs

Similarly, solution competition SPR experiments were used to evaluate the inhibitory effects of various GAGs on the binding of Bgp and DbpB-B31 to immobilize heparin. Bgp and DbpB-B31, at concentrations of 37.5 nM and 675 nM, respectively, were pre-mixed with 10 µM heparin, HS, CSA, DS, CSC, CSD, CSE, or KS. The mixtures were injected over a heparin-coated chip at 25 °C at a flow rate of 45 μL/min. The chip surface was regenerated after each cycle by injecting 45 μL of 2 M NaCl.

### 2.6. Inhibitory activity of sulfated glycans on the interaction between heparin and B. burgdorferi GAG-binding proteins

Solution competition SPR experiments were conducted to evaluate the inhibitory effects of marine-derived and related sulfated glycans on the binding of Bgp and DbpB-B31 to immobilized heparin. Bgp and DbpB-B31, at concentrations of 37.5 nM and 675 nM, respectively, were pre-mixed with IbSF, IbFucCS, PpFucCS, LvSF, HfSF, HfFucCS, MPS, PPS, RPI-28, and RS at a concentration of 10 µg/mL. The protein-glycan mixtures were injected over the heparin chip at a flow rate of 45 μL/min. After each cycle, the sensor surface was regenerated by injecting 45 μL of 2 M NaCl.

### 2.7. Statistical Analysis

Statistical analysis was performed using one-way ANOVA followed by Tukey’s multiple comparisons test. All statistical analyses were conducted using GraphPad Prism 8 software. Statistical significance was defined as follows: ns, *p* > 0.05; *, *p* < 0.05; **, *p* < 0.01; and ***, *p* < 0.001.

## 3. Results and discussion

### 3.1. Binding kinetics and affinity of B. burgdorferi GAG-binding adhesins with heparin

*B. burgdorferi* expresses several surface-associated adhesins that recognize GAGs, proteoglycans, and other extracellular-matrix components, thereby facilitating host-tissue colonization and dissemination. GAGs displayed on cell-surface proteoglycans or within the extracellular matrix provide potential attachment sites for the spirochete. Bgp, DbpA, and DbpB have been implicated in these interactions with host GAGs or extracellular-matrix ligands. Because of its high sulfation density and structural relationship to HS, heparin was used as a model ligand to characterize the GAG-binding kinetics and affinities of these proteins.

To compare the GAG-recognition properties of selected *B. burgdorferi* adhesins, the binding of Bgp, DbpA-A9, and DbpB-B31 to immobilized heparin was analyzed by SPR. All three proteins exhibited concentration-dependent binding responses. The sensorgrams were globally fitted to a 1:1 Langmuir binding model to determine the association rate constant (*ka*), dissociation rate constant (*kd*), and equilibrium dissociation constant (*K_D_*). The corresponding sensorgrams and binding kinetic parameters are shown in Fig. 2 and Table 1. Bgp exhibited the highest affinity for heparin, with a *K_D_* of 2.18 nM, followed by DbpB-B31 and DbpA-A9, with *K_D_* values of 36.4 and 88.4 nM, respectively. These results demonstrate distinct heparin-binding affinities among the three adhesins and support the involvement of sulfated GAGs in *B. burgdorferi* attachment and tissue colonization.

**Fig. 2.**
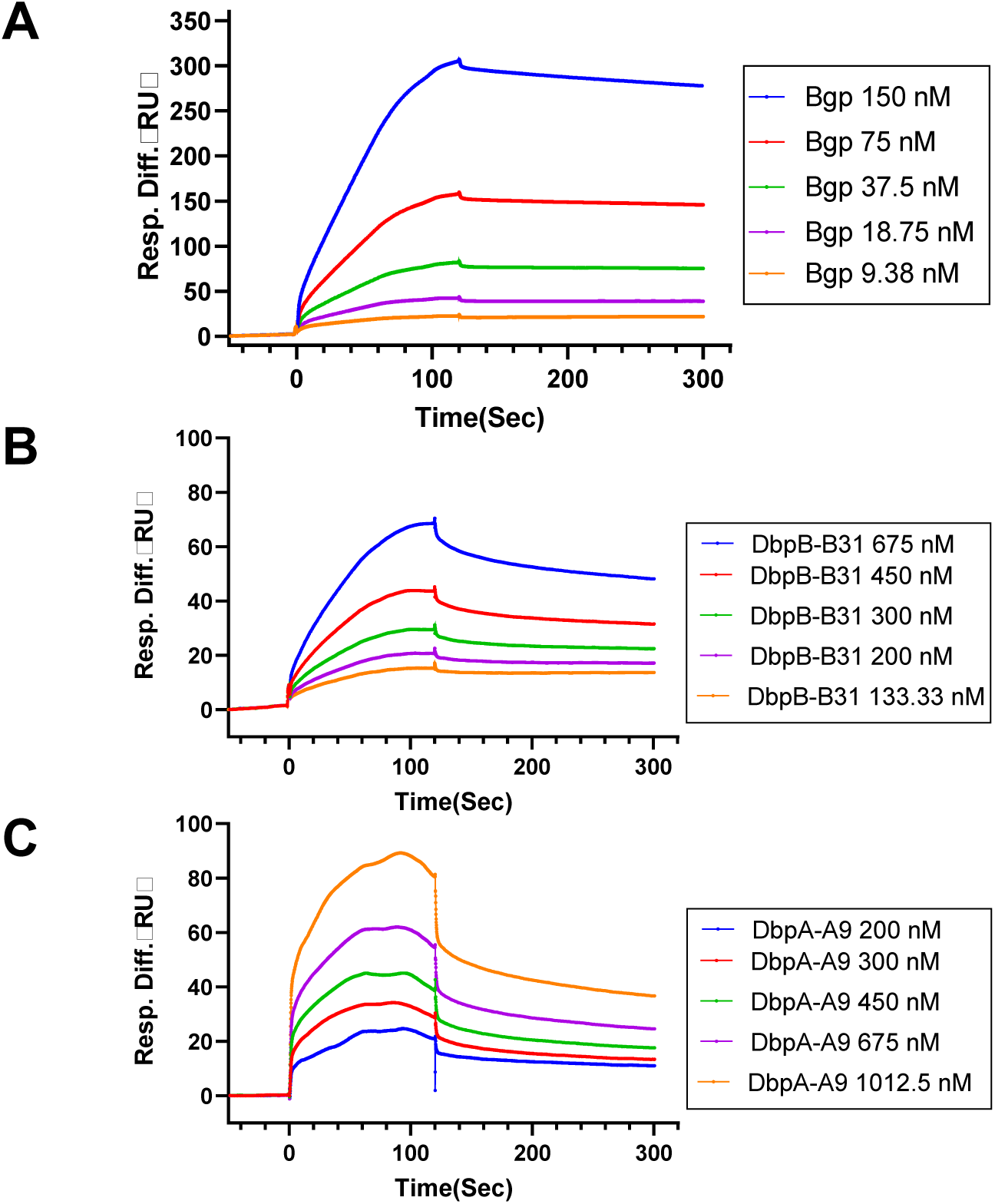
SPR kinetic analysis of Bgp, DbpA-A9, and DbpB-B31 binding to immobilized heparin. **(A)** SPR sensorgrams of Bgp binding to immobilized heparin at concentrations of 150, 75, 37.5, 18.75, and 9.38 nM. **(B)** SPR sensorgrams of DbpB-B31 binding to immobilized heparin at concentrations of 675, 450, 300, 200, and 133.33 nM. **(C)** SPR sensorgrams of DbpA-A9 binding to immobilized heparin at concentrations of 1012.5, 675, 450, 300, and 200 nM.

**Table 1.** Kinetic and affinity parameters for the binding of Bgp, DbpA-A9, and DbpB-B31 to immobilized heparin.

| | $k_a$ (M<sup>-1</sup>s<sup>-1</sup>) | $k_d$ (s<sup>-1</sup>) | $K_D$ (M) |
| --- | --- | --- | --- |
| Bgp | 1.83E+05 | 4.15E-04 | 2.18E-09 |
|  | (± 7.80E+04)* | (±2.46E-04)* | (±3.69E-10)** |
| DbpB-B31 | 4.53E+04 | 1.63E-03 | 3.64E-08 |
|  | (±6.51E+03)* | (±3.70E-04)* | (±9.03E-09)** |
| DbpA-A9 | 2.86E+04 | 2.50E-03 | 8.84E-08 |
|  | (±3.14E+03)* | (±3.66E-04)* | (±1.35E-08)** |

\*The data with (±) in parentheses are standard deviations (SD) from global fitting of five injections (with different concentrations of proteins). \*\*The data with (±) in parentheses are standard deviations (SD) from triplicate measurements.

The higher heparin-binding affinity of Bgp than DbpA-A9 and DbpB-B31 suggests that these adhesins differ in their recognition of sulfated GAG structures. Previous studies have similarly demonstrated distinct GAG-binding specificities for DbpA and DbpB and identified different GAG-binding-site architectures in these two proteins (Feng & Wang, 2015; Fischer et al., 2003). These molecular differences may support complementary roles in cell-specific attachment and tissue colonization during infection (Schlachter, Seshu, Lin, Norris, & Parveen, 2018). Bgp binds heparin-related GAGs, and its loss impairs spirochetal adherence and tissue colonization, although it is not essential for infection (Schlachter et al., 2018). Thus, the nanomolar affinities observed here further support the biological relevance of GAG-mediated interactions in *B. burgdorferi* attachment. However, because heparin is generally more highly sulfated than host HS, further studies using physiologically relevant GAG structures are needed to define the native ligand preferences of these adhesins.

Based on their distinct heparin-binding characteristics, the secreted surface and GAG-binding protein Bgp and another surface-associated adhesin DbpB-B31 were selected as representative targets for subsequent competitive inhibition assays (Parveen & Leong, 2000). This selection strategy accounts for host interactions mediated by a pathogen, thereby providing an experimental basis for evaluating the ability of sulfated polysaccharides to interfere with these two types of interactions.

### 3.2. Inhibitory activity of heparin oligosaccharides and desulfated heparins on protein-heparin interactions

HSPGs are widely distributed on host-cell surfaces and within the extracellular matrix and can serve as potential binding sites for *B. burgdorferi* attachment and tissue colonization. Given that *B. burgdorferi* expresses multiple GAG-binding proteins that differ in their structure, and biological functions, individual proteins may have distinct requirements for glycan chain length and sulfation pattern. Based on the preceding binding analyses, we selected two representative proteins for subsequent competition assays with sulfated polysaccharides: Bgp, a multifunctional GAG-binding protein that exhibited the highest affinity for heparin and can be shed into the extracellular environment under adverse growth conditions, and DbpB-B31, a surface-associated adhesin with a distinct GAG-binding architecture. Evaluating these functionally and structurally different proteins enabled us to compare their glycan-recognition preferences and assess the ability of candidate polysaccharides to competitively inhibit *B. burgdorferi* adhesion-GAG interactions.

To determine how heparin chain length and sulfation pattern affect the binding of Bgp and DbpB-B31, we performed solution-phase competitive SPR assays using size-defined heparin oligosaccharides and selectively desulfated heparin derivatives. Soluble competitors were premixed with the proteins before injection over the heparin-immobilized sensor chip. Occupancy of the protein-binding sites by soluble competitors reduced subsequent binding signal (RU) to the immobilized heparin. Therefore, lower relative SPR responses indicated greater competitive potency.

The heparin oligosaccharide competition assays revealed distinct chain-length requirements for Bgp and DbpB-B31 (Fig. 3). Bgp showed little apparent dependence on chain length across the dp4-dp20 range, although all tested oligosaccharides showed weaker competitive activity than unfractionated heparin. In contrast, DbpB-B31 exhibited a marked chain-length dependence, with longer oligosaccharides showing progressively greater competitive activity and a notable increase in inhibition at dp12 and above. However, the inhibitory effects of these oligosaccharides remained weaker than those of unfractionated heparin. This preference may indicate that DbpB-B31 requires an extended glycan-binding interface or engages multiple basic regions along the heparin chain. The greater inhibitory activity of unfractionated heparin suggests that the higher-molecular-weight polymer provides a more effective presentation of sulfated motifs under the tested conditions. This protein-specific chain-length dependence is consistent with earlier evidence that longer heparin chains more effectively inhibit *B. burgdorferi* attachment to mammalian cells (Leong et al., 1998). The preference of DbpB-B31 for longer oligosaccharides may reflect its extended GAG-binding surface, including a lysine-rich C-terminal region known to contribute substantially to GAG recognition (Feng & Wang, 2015). The limited chain-length dependence of Bgp suggests that shorter sulfated sequences may be sufficient to occupy its principal GAG-binding site. In fact, Bgp possesses a motif matching significantly to previously described heparin-binding consensus sequence in other known human and microbial proteins (Fig. 3E). Such domain is not apparent in the previously proposed heparin-binding peptides of DbpB (Feng & Wang, 2015). Together, these findings indicate that Bgp and DbpB-B31 recognize heparin through distinct binding mechanisms, which may support their complementary functions in host attachment and tissue colonization.

**Fig. 3.**
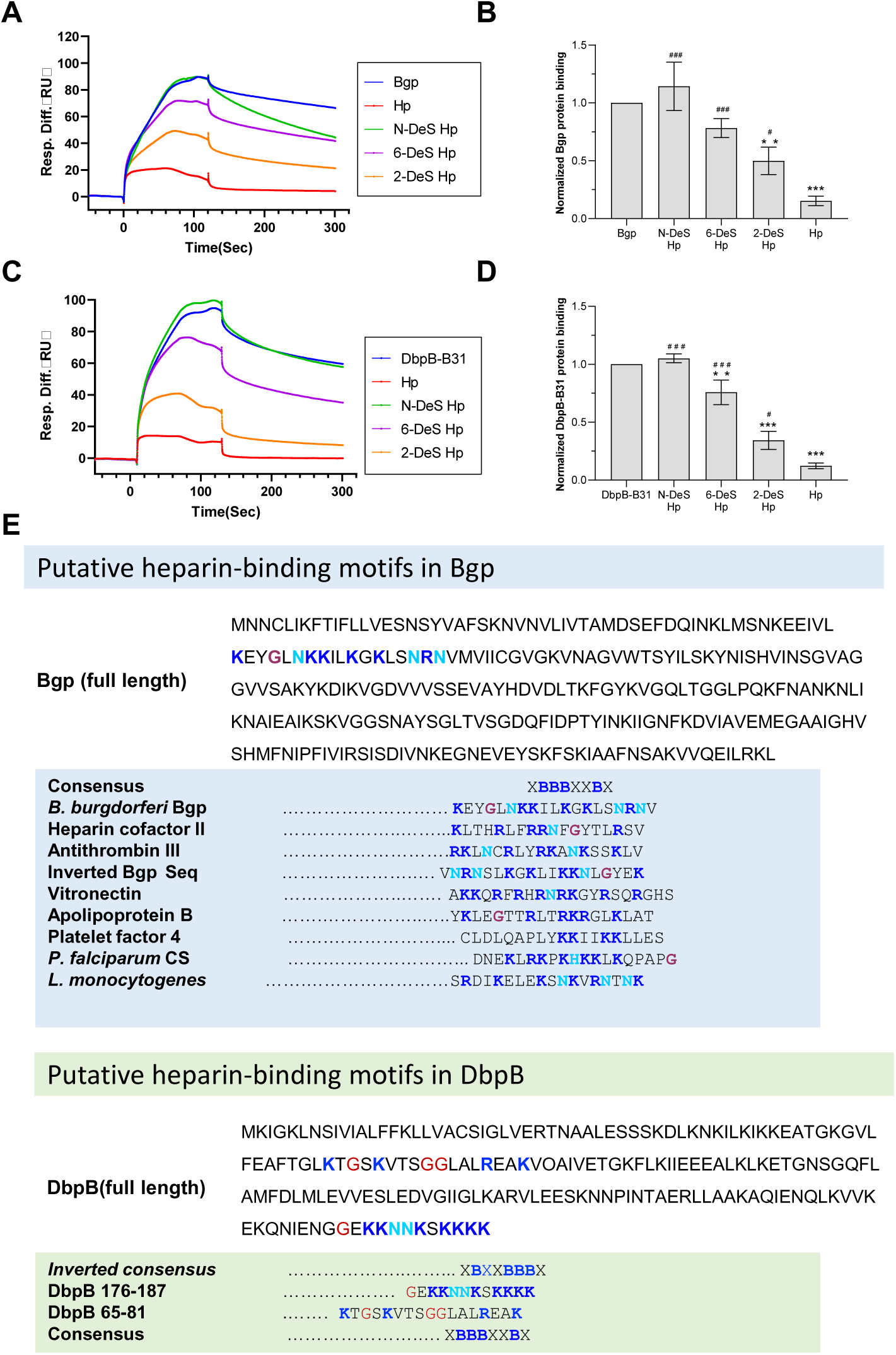
Effects of heparin chain length on the binding of Bgp and DbpB-B31 to immobilized heparin. **(A)** SPR sensorgrams of the Bgp-heparin interaction in competition with soluble heparin and heparin oligosaccharides. Bgp at 37.5 nM was pre-mixed with 10 μM heparin or the indicated heparin oligosaccharide. **(B)** Bar graph showing the normalized binding of Bgp to surface-immobilized heparin in the presence of the indicated competitors. **(C)** SPR sensorgrams of the DbpB-B31-heparin interaction in competition with soluble heparin and heparin oligosaccharides. DbpB-B31 at 675 nM was pre-mixed with 10 μM heparin or the indicated heparin oligosaccharide. **(D)** Bar graph showing the normalized binding of DbpB-B31 to surface-immobilized heparin in the presence of the indicated competitors. **(E)** Sequence comparison of putative heparin-binding motifs in Bgp and DbpB-B31 with representative basic heparin-binding sequences. Conserved Lys/Arg-rich residues are highlighted. Bar graphs represent the mean ± SD from three replicate measurements.

Heparin sulfation patterns are critical determinants of heparin–protein interactions. Competition assays using selectively desulfated heparins showed that *N*-desulfation produced the greatest loss of inhibitory activity against both Bgp and DbpB-B31, identifying *N*-sulfate groups as major determinants of heparin recognition (Fig. 4). 6-*O*-desulfation also substantially decreased competitive activity, supporting an important contribution of 6-*O*-sulfate groups. In contrast, 2-*O*-desulfation had the smallest effect, indicating a comparatively limited role for 2-*O*-sulfate groups. Thus, the apparent contributions of the sulfate positions to the binding of both proteins followed the order *N*-sulfation > 6-*O*-sulfation > 2-*O-* sulfation. These findings are consistent with earlier studies showing that GAG sulfation is essential for efficient *B. burgdorferi* attachment to host cells. *N*-desulfated and extensively *O*-desulfated heparin derivatives were previously reported to lose their ability to inhibit spirochetal attachment, demonstrating the importance of sulfate-dependent electrostatic interactions (Isaacs, 1994). In addition, the inhibitory potency of heparin and heparan sulfate fractions was shown to increase with their overall charge density (Leong et al., 1998). Our results extend these observations by distinguishing the contributions of individual sulfate positions and identifying *N*-sulfation and 6-*O*-sulfation as the principal determinants of Bgp and DbpB-B31 binding. The similar sulfation requirements of the two proteins suggest a shared preference for *N*- and 6-*O*-sulfated domains, whereas the relatively small contribution of 2-*O*-sulfation indicates that overall negative charge alone does not fully govern recognition.

**Fig. 4.**
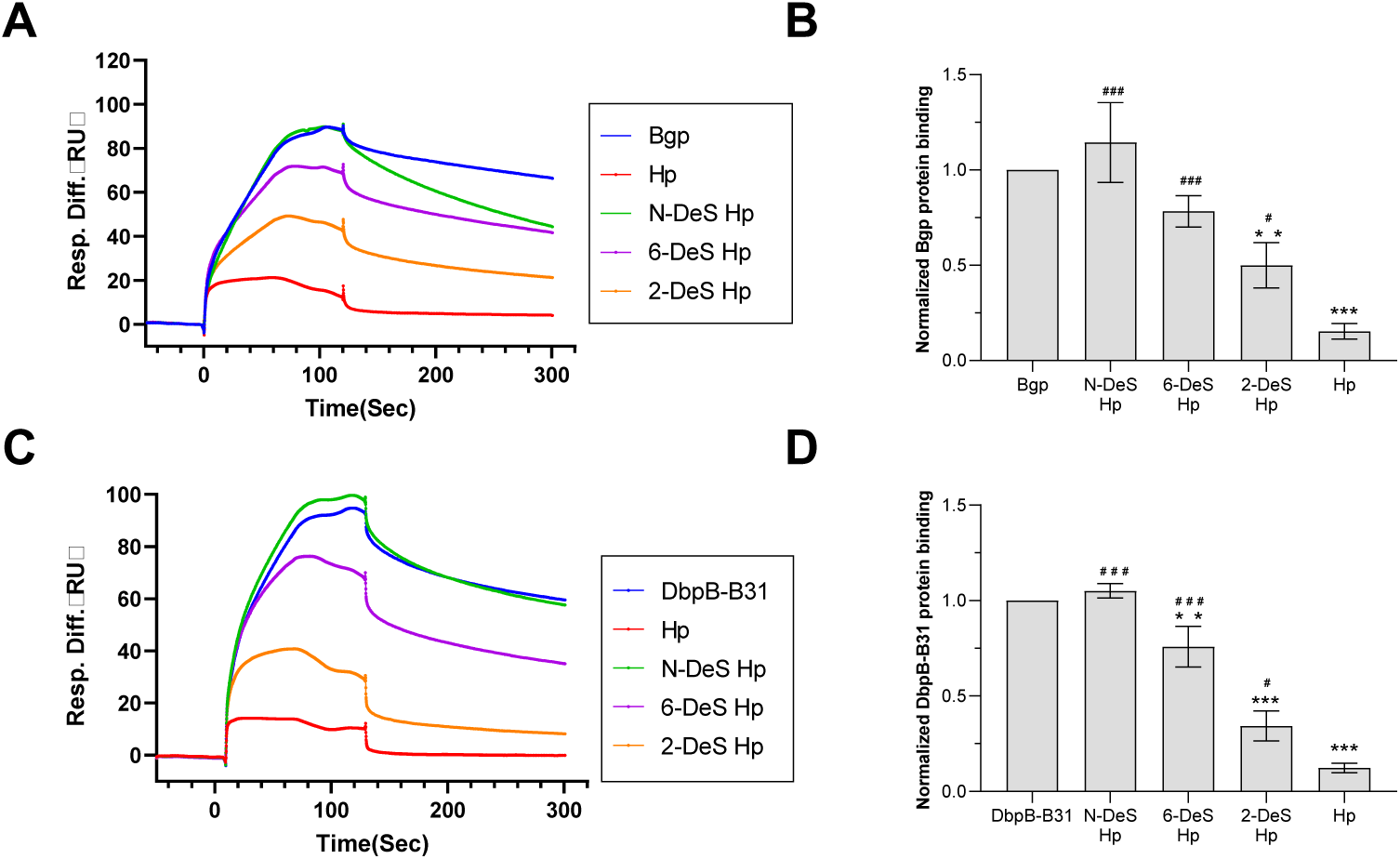
Effects of sulfation of heparin on Bgp and DbpB-B31 binding to immobilized heparin. (A) SPR sensorgrams of the Bgp-heparin interaction in competition with different selective desulfated heparins. Bgp at 37.5 nM was pre-mixed with 10 μM of the indicated soluble competitor. (B) Bar graph showing the normalized binding of Bgp to surface-immobilized heparin in the presence of the indicated heparin derivatives. (C) SPR sensorgrams of the DbpB-B31-heparin interaction in competition with different selective desulfated heparins. DbpB-B31 at 675 nM was pre-mixed with 10 μM of the indicated soluble competitor. (D) Bar graph showing the normalized binding of DbpB-B31 to surface-immobilized heparin in the presence of heparin derivatives. Bar graphs represent the mean ± SD from three replicate measurements.

### 3.3. Inhibitory activity of various GAGs on the heparin binding of Bgp and DbpB-B31

Structurally diverse GAGs are present on host cell surfaces and within the extracellular matrix and may serve as ligands for *B. burgdorferi* adhesins. Because GAG composition and sulfation patterns vary among tissues, defining the GAG selectivity of Bgp and DbpB-B31 may provide insight into their differential recognition of host tissues. To characterize this selectivity, we used competitive SPR assays to evaluate heparin, HS, CSA, CSC, CSD, CSE, DS and KS as inhibitors of protein binding to immobilized heparin.

The competition SPR assays showed that the tested GAGs inhibited the binding of Bgp and DbpB-B31 to immobilized heparin to different levels (Fig. 5). CSE produced the greatest inhibition of both proteins under the tested conditions. Its high sulfation density, arising primarily from 4-*O*- and 6-*O*-disulfated *N*-acetylgalactosamine residues, may create a spatial charge distribution favorable for recognition by both adhesins. The inhibitory activities of several other GAGs suggest that Bgp and DbpB-B31 can recognize sulfated glycans with diverse backbone structures. Notably, KS showed slightly greater inhibition of Bgp than DS. KS contains abundant 6-*O*-sulfated residues, whereas DS is characterized primarily by iduronic acid–*N*-acetylgalactosamine 4-*O*-sulfate repeating units. This difference is consistent with the important contribution of 6-*O*-sulfation identified using selectively desulfated heparins. However, the strong activity of CSE indicates that GAG recognition is not determined by a single sulfate position but likely reflects the combined effects of backbone composition, sulfation density, and spatial sulfate distribution. Collectively, these results demonstrate that Bgp and DbpB-B31 have broad but nonidentical GAG-recognition profiles, with a preference for highly sulfated structures. These findings support the subsequent evaluation of non-heparin and marine-derived sulfated polysaccharides as competitive inhibitors.

**Fig. 5.**
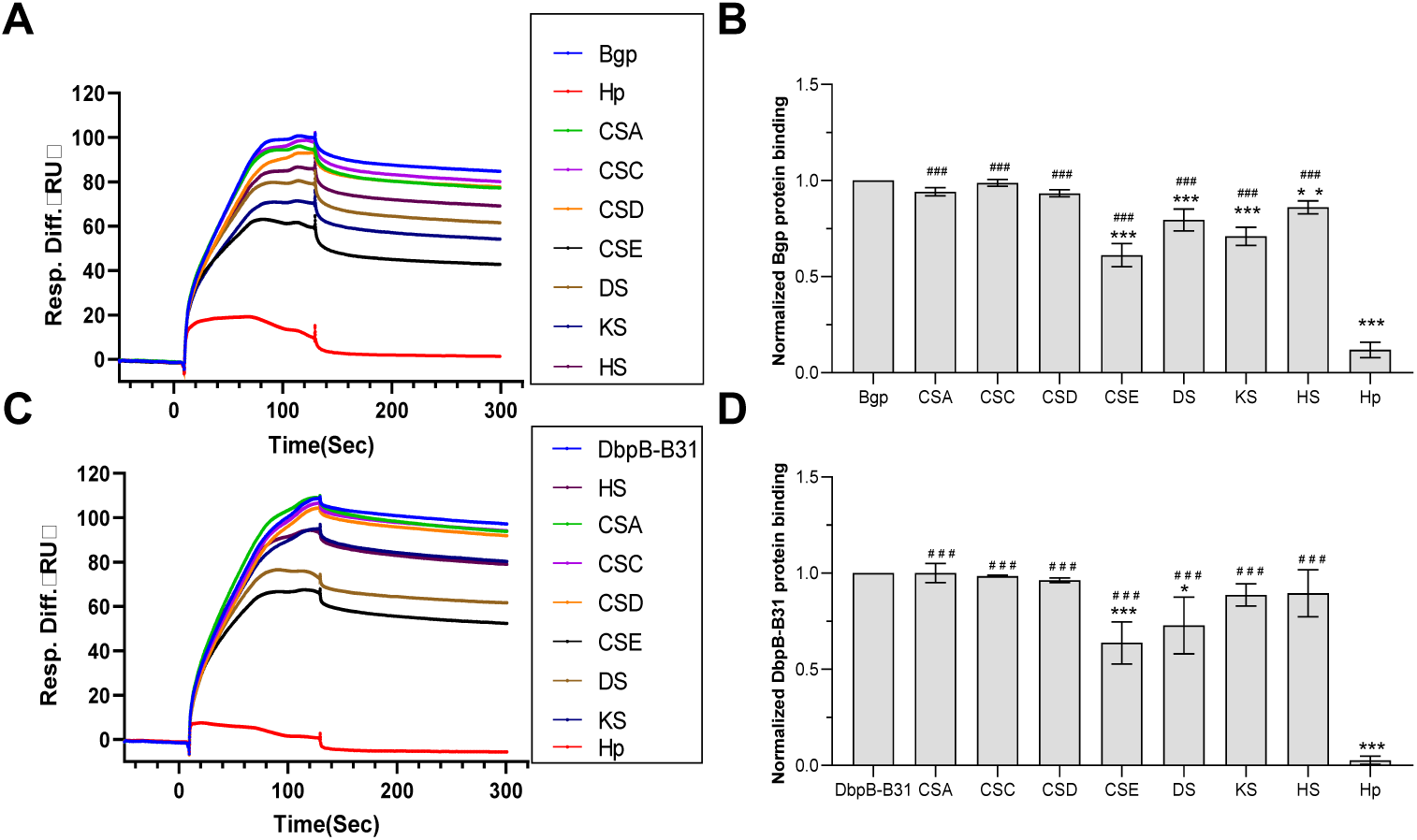
Inhibition of the binding of Bgp and DbpB-B31 to immobilized heparin by different GAGs. (A) SPR sensorgrams of the Bgp-heparin interaction in competition with different GAGs. Bgp at 37.5 nM was pre-mixed with 10 μM of the indicated soluble competitor. (B) Bar graph showing the normalized binding of Bgp to surface-immobilized heparin in the presence of the indicated heparin derivatives. (C) SPR sensorgrams of the DbpB-B31-heparin interaction in competition with different GAGs. DbpB-B31 at 675 nM was pre-mixed with 10 μM of the indicated soluble competitor. (D) Bar graph showing the normalized binding of DbpB-B31 to surface-immobilized heparin in the presence of different GAGs. Bar graphs represent the mean ± SD from three replicate measurements.

The broad but differential GAG-recognition profiles observed here are consistent with previous reports that *B. burgdorferi* adhesins recognize multiple sulfated GAGs. Bgp has been shown to bind heparin and dermatan sulfate, whereas DbpA and DbpB mediate GAG-dependent attachment with distinct ligand and cell-type preferences (Fischer et al., 2003; Parveen & Leong, 2000). Earlier whole-cell studies also demonstrated that different classes of proteoglycans contribute to spirochetal attachment to epithelial, endothelial, and neural cells (Leong et al., 1998). Our identification of CSE as the strongest competitor for both Bgp and DbpB-B31 extends these observations by highlighting the importance of high local sulfation density rather than a heparin/HS-specific backbone. These results suggest that tissue-specific differences in GAG structure may influence adhesin-mediated colonization and that highly sulfated non-heparin glycans may serve as effective competitive inhibitors.

### 3.4. Inhibitory activity of PPS and MPS on protein-heparin interactions

Pentosan polysulfate (PPS) and mucopolysaccharide polysulfate (MPS) are highly sulfated polysaccharides with high negative charge densities (Fig.1). We hypothesized that these compounds could competitively inhibit the binding of Bgp and DbpB-B31 to heparin by occupying their GAG-binding sites. To evaluate this possibility, competitive SPR assays were used to compare the inhibitory effects of PPS and MPS on the binding of Bgp and DbpB-B31 to immobilized heparin (Fig. 6).

**Fig. 6.**
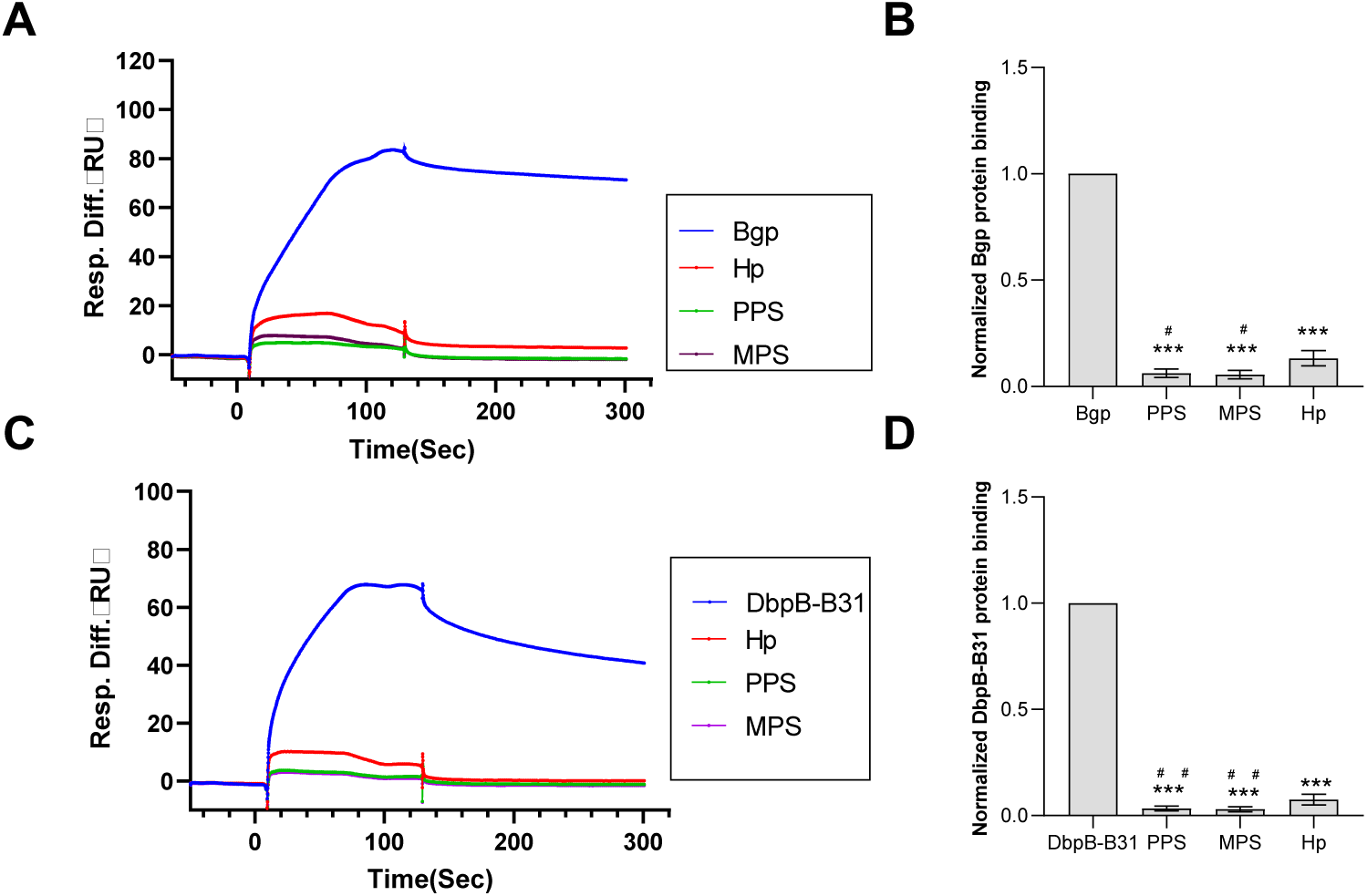
Inhibition of Bgp-/ DbpB-B31-heparin interactions by PPS and MPS. (A) SPR sensorgrams of the Bgp-heparin interaction in competition with PPS and MPS. Bgp at 37.5 nM was pre-mixed with 10 μg/mL of sulfated glycans. (B) Bar graph showing the normalized binding of Bgp to surface-immobilized heparin in the presence of sulfated glycans. (C) SPR sensorgrams of the DbpB-B31–heparin interaction in competition with PPS and MPS. DbpB-B31 at 675 nM was pre-mixed with 10 μg/mL of sulfated glycans. (D) Bar graph showing the normalized binding of DbpB-B31 to surface-immobilized heparin in the presence of sulfated glycans. Bar graphs represent the mean ± SD from three replicate measurements.

For DbpB-B31, PPS exhibited the strongest competitive activity among all polysaccharides tested, with MPS ranking second, indicating efficient recognition of both highly sulfated glycans by this adhesin. For Bgp, both PPS and MPS inhibited binding to immobilized heparin, but their activities were not substantially greater than that of heparin under the tested conditions. Thus, PPS and MPS displayed protein-dependent inhibition profiles, with PPS showing particularly strong activity against DbpB-B31. These findings identified PPS as the leading candidate for subsequent concentration-response, IC_50_ and cell-adhesion studies, with MPS included as a structurally distinct comparator for further structure-activity analysis. PPS and MPS are highly sulfated GAG mimetics previously shown to inhibit interactions between heparin and heparin-binding pathogen proteins (F. Zhang et al., 2022). Their inhibition of Bgp and DbpB-B31 therefore supports the broader concept that soluble sulfated polysaccharides can function as competitive decoys for *B. burgdorferi* GAG-binding adhesins (Lin, Li, Zhang, & Linhardt, 2017).

### 3.5. Inhibitory activity of marine invertebrate-derived sulfated glycans on protein-heparin interactions

Marine invertebrate-derived sulfated polysaccharides possess structurally diverse glycan backbones, branching patterns, and sulfate distributions that may differentially affect their recognition by B. burgdorferi GAG-binding proteins. To evaluate how these structural features influence competitive activity, we examined the effects of the sea cucumber-derived polysaccharides IbSF, IbFucCS, PpFucCS, HfSF, and HfFucCS and the sea urchin-derived polysaccharide LvSF on the binding of Bgp and DbpB-B31 to immobilized heparin (Fig.7).

**Fig. 7.**
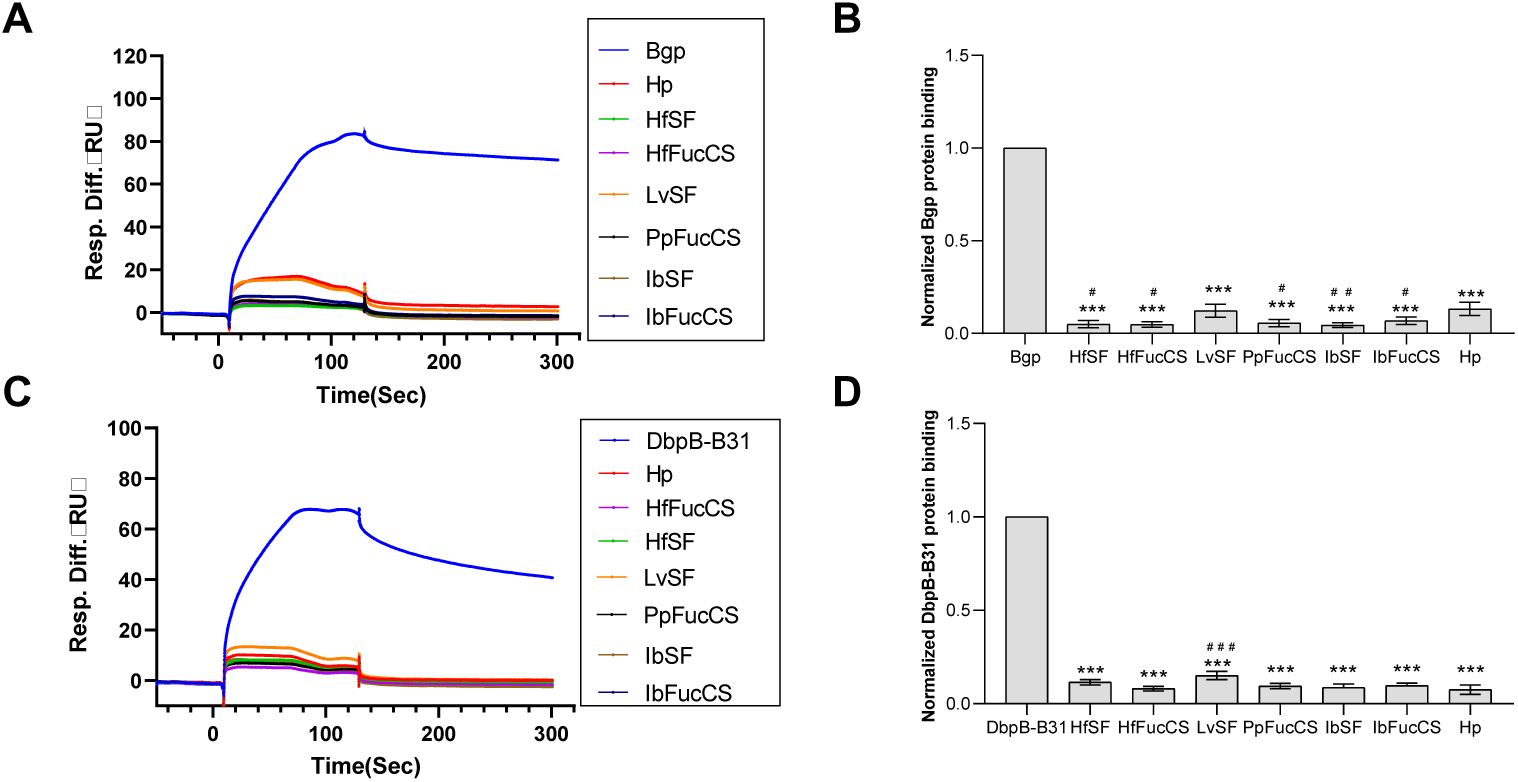
Inhibition of Bgp-/DbpB-B31-heparin interactions by marine invertebrate-derived sulfated glycans. (A) SPR sensorgrams of the Bgp-heparin interaction in competition with different marine sulfated glycans. Bgp (37.5 nM) was pre-mixed with each sulfated glycan or soluble heparin at 10 μg/mL. (B) Bar graph showing the normalized binding of Bgp to surface-immobilized heparin in the presence of different sulfated glycans. (C) SPR sensorgrams of the DbpB-B31–heparin interaction in competition with different marine sulfated glycans. DbpB-B31 (675 nM) was pre-mixed with each sulfated glycan at 10 μg/mL. (D) Bar graph showing the normalized binding of DbpB-B31 to surface-immobilized heparin in the presence of different sulfated glycans. Bar graphs represent the mean ± standard deviation from triplicate experiments.

Sulfated polysaccharides from different marine sources exhibited distinct inhibitory activities against Bgp and DbpB-B31. For Bgp, fucosylated chondroitin sulfate (FucCS) preparations generally showed greater competitive activity than sulfated fucans (SFs), with HfFucCS producing the largest reduction in binding. Because FucCS and SF isolated from the same organisms differ in their glycan backbones, branching patterns, and sulfate-group distributions, these results indicate that Bgp recognition is not governed solely by overall negative charge or degree of sulfation. Instead, structural features such as backbone composition, chain conformation, fucosyl branching, and the spatial presentation of sulfate groups may contribute to binding. DbpB-B31 displayed a different inhibition profile, further indicating that Bgp and DbpB-B31 possess distinct preferences for sulfated-polysaccharide structures. Collectively, these results demonstrate that marine invertebrate-derived sulfated polysaccharides can competitively inhibit the binding of both proteins to heparin and identify HfFucCS as a particularly potent inhibitor of Bgp.

Previous studies have shown that marine FucCSs and SFs can act as GAG mimetics and competitively inhibit interactions between pathogen proteins and heparin or heparan sulfate (Dwivedi et al., 2021; He et al., 2023). Their inhibitory activities vary substantially among glycans and protein targets, indicating that activity depends not only on sulfate density but also on molecular weight, backbone structure, fucosyl branching, sulfation pattern, and chain conformation. The strong inhibition of Bgp by HfFucCS observed here is consistent with this structure-dependent recognition. Moreover, the different inhibition profiles of Bgp and DbpB-B31 suggest that the structural complementarity between each glycan and the adhesin-binding surface determines competitive potency. These findings extend the potential application of marine sulfated glycans from antiviral targets to bacterial anti-adhesion strategies.

### 3.6. Inhibitory activity of algal-derived sulfated glycans on protein-heparin

Algal-derived sulfated glycans possess backbone structures and sulfation patterns distinct from those of heparin, chondroitin sulfate, and sea cucumber-derived sulfated polysaccharides. To assess their competitive activities against Bgp and DbpB-B31, we performed solution-phase competitive SPR assays using the brown algal polysaccharide (fucoidan) RPI-28 and the green algal polysaccharide rhamnan sulfate (RS). Both glycans reduced the binding of Bgp and DbpB-B31 to immobilized heparin, indicating that they are recognized by these proteins and can compete with heparin for their GAG-binding sites (Fig. 8).

**Fig. 8.**
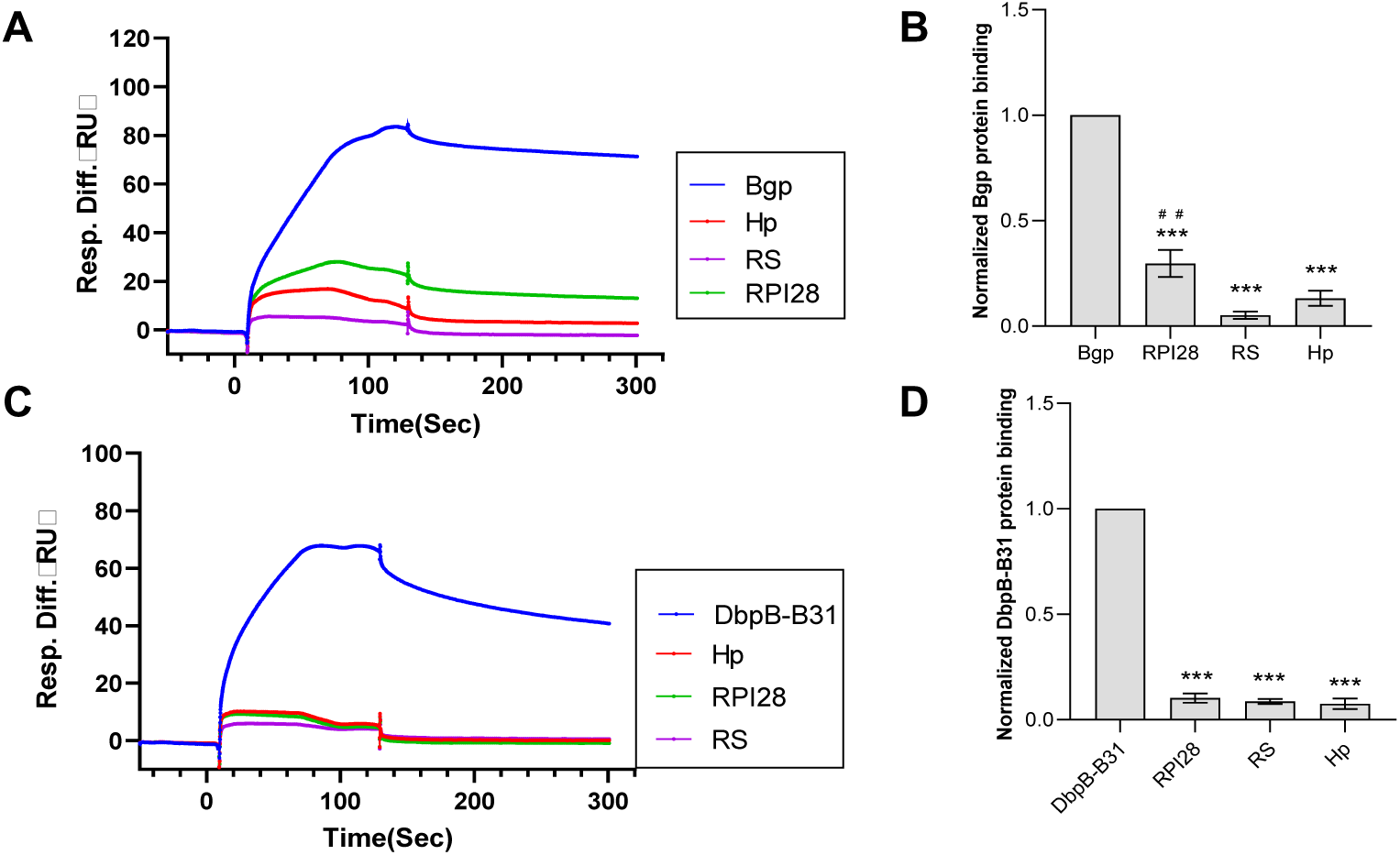
Inhibition of Bgp and DbpB-B31 binding to immobilized heparin by algal-derived sulfated glycans. (A) SPR sensorgrams of the Bgp-heparin interaction in competition with RPI-28, RS, or soluble heparin. Bgp at 37.5 nM was pre-mixed with 10 μg/mL of sulfated glycans. (B) Bar graph showing the normalized binding of Bgp to surface-immobilized heparin in the presence of sulfated glycans. (C) SPR sensorgrams of the DbpB-B31-heparin interaction in competition with sulfated glycans. DbpB-B31 at 675 nM was pre-mixed with 10 μg/mL of sulfated glycans. (D) Bar graph showing the normalized binding of DbpB-B31 to surface-immobilized heparin in the presence of sulfated glycans. Bar graphs represent the mean ± SD from three replicate measurements.

Both RPI-28 and RS competitively inhibited the binding of Bgp and DbpB-B31 to immobilized heparin, although their inhibitory activities differed between the two proteins. These protein-dependent inhibition profiles suggest that activity is influenced by structural features beyond overall sulfation, including glycan backbone composition, sulfation pattern, and spatial charge distribution. RS exhibited particularly strong inhibition of Bgp and was therefore selected for subsequent concentration-response and IC_50_ analyses as a potential non-animal-derived inhibitor of GAG-adhesin interactions.

The inhibitory activity of these algal glycans indicates that a heparin-like saccharide backbone is not required for recognition by Bgp or DbpB-B31. RS, which is primarily composed of sulfated α-L-rhamnose residues, has previously been shown to bind the SARS-CoV-2 spike protein and potently inhibit its interaction with immobilized heparin (Song et al., 2021). Fucoidans and RS from different algal species also exhibit structure-dependent inhibition of protein-HS interactions, with activity influenced by molecular weight, backbone composition, and sulfation pattern (Song et al., 2024). The strong inhibition of Bgp by RS therefore supports its function as a non-animal-derived GAG mimetic. The differing activities of RPI-28 and RS against Bgp and DbpB-B31 further indicate that their potency depends on structural complementarity with individual adhesin-binding surfaces rather than negative charge alone.

### 3.7. Concentration-dependent inhibitory effects of sulfated glycans

Single-concentration competition assays demonstrated that several natural and semisynthetic sulfated glycans inhibited the binding of Bgp and DbpB-B31 to immobilized heparin. To quantitatively compare their inhibitory potencies, we performed concentration-response competition assays with heparin, IbFucCS, PpFucCS, HfFucCS, RS, and PPS. The residual binding response at each competitor concentration was normalized to that of a protein-only control, and the resulting concentration–response curves were fitted by nonlinear regression to determine the apparent IC_50_ values (Fig. 9 and Table 2).

**Fig. 9.**
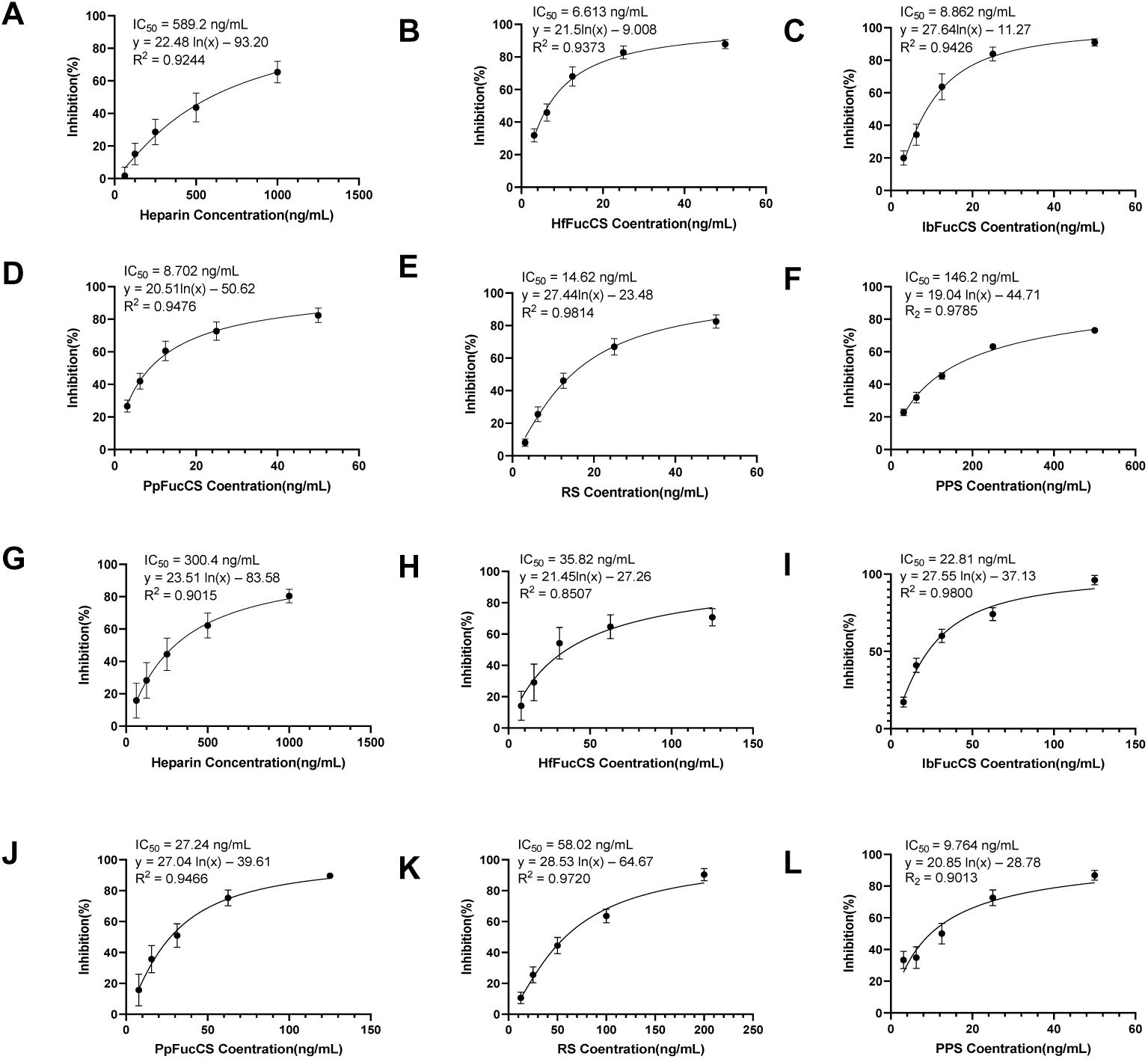
Concentration-dependent inhibition of Bgp–heparin and DbpB-B31–heparin interactions by sulfated glycans. The inhibitory effects of heparin, HfFucCS, IbFucCS, PpFucCS, rhamnan sulfate (RS), and pentosan polysulfate (PPS) on the binding of Bgp and DbpB-B31 to surface-immobilized heparin were evaluated using solution competition SPR assays. Panels A–F show the concentration-dependent inhibition of Bgp by (A) heparin, (B) HfFucCS, (C) IbFucCS, (D) PpFucCS, (E) RS, and (F) PPS. Panels G–L show the corresponding inhibition of DbpB-B31 by (G) heparin, (H) HfFucCS, (I) IbFucCS, (J) PpFucCS, (K) RS, and (L) PPS. Each sulfated glycan was pre-mixed with the corresponding protein before injection over the heparin-coated sensor surface. The binding response obtained in the absence of a soluble competitor was used as the control, and inhibition was calculated as the percentage reduction in binding response relative to this control. Solid lines represent logarithmic regression fits used to estimate the apparent half-maximal inhibitory concentrations (IC_50_); the fitted equations, IC_50_ values, and coefficients of determination (R²) are shown in each panel. Data are presented as mean ± SD from 3 replicate measurements.

**Table 2.** Apparent IC_50_ values of sulfated glycans for inhibition of Bgp and DbpB-B31 binding to immobilized heparin.

|  | Heparin | HfFucCS | IbFucCS | PpFucCS | RS | PPS |
| --- | --- | --- | --- | --- | --- | --- |
| Bgp (ng/mL) | 589.2 | 6.6 | 8.9 | 8.7 | 14.6 | 146 |
| DbpB-B31 (ng/mL) | 300.4 | 35.8 | 22.8 | 27.2 | 58.0 | 9.8 |

All sulfated glycans that achieved sufficient inhibition over the tested concentration range displayed concentration-dependent responses suitable for nonlinear regression analysis, although their apparent IC_50_ values varied widely. For Bgp, FucCSs generally exhibited strong inhibitory activity, with HfFucCS showing the lowest IC₅₀ value of 6.6 ng/mL. In comparison, unfractionated heparin had an IC_50_ of 589.2 ng/mL. On a mass-concentration basis, HfFucCS was therefore nearly 90-fold more potent than heparin under these experimental conditions. IbFucCS and PpFucCS also demonstrated clear concentration-dependent inhibition but were less potent than HfFucCS. The inhibitory activities observed for the SFs (IbSF, LvSF, and HfSF) and for RPI-28, RS, MPS, and PPS further demonstrate that Bgp recognition is not restricted to heparin- or HS-like glycan backbones.

In contrast to Bgp, DbpB-B31 was most strongly inhibited by PPS, which produced a concentration-dependent reduction in binding to immobilized heparin and yielded the lowest apparent IC_50_ among the tested polysaccharides (9.8 ng/mL). MPS was less potent than PPS but more potent than several naturally derived marine glycans. FucCSs, SFs, RPI-28, and RS also produced measurable concentration-dependent inhibition; however, their potency ranking differed markedly from that observed for Bgp. These protein-specific inhibition profiles suggest that Bgp and DbpB-B31 differ in their polysaccharide-binding interfaces and structural requirements.

Overall, the IC_50_ analyses identified HfFucCS as the most potent inhibitor of Bgp and PPS as the most potent inhibitor of DbpB-B31. Because these polysaccharides differ substantially in backbone structure and molecular weight, their IC_50_ values represent apparent potencies based on mass concentration and should be interpreted accordingly. The moderate-to-strong activities of several additional polysaccharides provide a foundation for further structure-activity relationship studies. Moreover, the complementary selectivity of HfFucCS and PPS toward two GAG-binding proteins with different localization and functional properties supports evaluating their individual and combined effects in subsequent adhesion assays.

## 4. CONCLUSION

This study defined the heparin-binding properties of three *B. burgdorferi* GAG-binding proteins and evaluated sulfated polysaccharides as competitive inhibitors. SPR analysis showed that Bgp, DbpA-A9, and DbpB-B31 bound heparin with distinct affinities, with Bgp exhibiting the highest affinity. Competition assays identified *N*-sulfation and 6-*O*-sulfation as major binding determinants. Bgp showed little dependence on heparin oligosaccharide chain length over dp4-dp20, whereas DbpB-B31 showed greater inhibition with longer oligosaccharides, particularly dp12 and above; however, all tested oligosaccharides were fewer effective competitors than unfractionated heparin.

Conventional GAGs and marine-and algal-derived sulfated polysaccharides inhibited both proteins in a structure and protein dependent manner. CSE showed strong activity against both proteins, while HfFucCS was the most potent inhibitor of Bgp, with an apparent IC_50_ of 6.6 ng/mL, nearly 90-fold more potent than heparin on a mass-concentration basis. PPS was the most potent inhibitor of DbpB-B31, followed by MPS. These findings demonstrate that glycan backbone, chain length, and sulfate-group distribution collectively govern recognition by *B. burgdorferi* adhesins. HfFucCS and PPS represent promising leads for developing glycan-based anti-adhesion agents that could limit the initial attachment and tissue colonization of B. burgdorferi. Potential applications include topical prophylactic formulations for administration at tick-bite sites, coatings or dressings that reduce local spirochetal attachment, and combination inhibitors targeting multiple GAG-binding adhesins. These glycans may also serve as molecular probes for mapping adhesin specificity and guiding the design of smaller, structurally defined GAG mimetics. Further cell-based and animal studies are required to evaluate their efficacy, toxicity, and suitability for local administration. This study establishes a structural basis for targeting *B. burgdorferi*-host GAG interactions and provides a foundation for developing nonbactericidal, glycan-based strategies to prevent Lyme disease. The findings define distinct inhibition profiles of structurally diverse sulfated polysaccharides in purified adhesion-heparin systems.

## CRediT authorship contribution statement

**Changkai Bu:** Analysis, Data Curation, original draft preparation**. Ke Xia:** Supervision, Review and Editing**. Carly Fernandes:** Protein preparation**. Yi-Pin Lin:** Supervision, Review and Editing**. Nikhat Parveen:** Resource, Review and Editing. **Vitor H. Pomin:** Resource, Review and Editing. **Jonathan S. Dordick:** Supervision, Review and Editing. **Lianli Chi:** Supervision, Review and Editing. **Fuming Zhang:** Conceptualization, Supervision, Original draft preparation, Funding acquisition.

## Declaration of Competing Interest

The authors declare that they have no known competing financial interests or personal relationships that could have appeared to influence the work reported in this paper.

## Acknowledgment

This research was partially funded by grants from the National Institutes of Health: S10OD028523 and S10OD032168 (F. Z.).

## Data availability

Data will be made available on request.

## TABLES

**Table 1.** MW parameters, distribution profiles, and anti-factor Xa activities of heparins derived from different animal sources.

**Table 2.** MW parameters, distribution profiles, and anti-factor Xa activities of LMWHs derived from different animal sources.

## FIGURE LEGENDS

**Figure 1.** Source-dependent differences in heparin-induced platelet activation. Representative flow cytometry scatter plots (top and bottom left) display the expression of activation markers (CD62P and CD63) following treatment with PMH, OMH, or BLH, with the right quadrants (Q1-LR) designating activated platelet populations. The accompanying bar graph (bottom right) presents the quantitative analysis of platelet activation rates (%). Data are presented as mean ± SD (n = 3). Statistical analysis was performed using one-way ANOVA, **P < 0.01 vs PMH, ***P < 0.001 vs PMH, and ^#^P < 0.05 vs OMH.

**Figure 2.** Source-dependent recognition of HIT antibodies by PF4–heparin complexes.(A) Schematic of the competitive ELISA protocol. (B) Competitive ELISA results demonstrating pronounced source-dependent variations in antibody recognition, expressed as absolute OD values (top) and relative antibody binding percentages (bottom). Data are presented as mean ± SD (n = 3). Statistical analysis was performed using one-way ANOVA, **P < 0.01 vs PMH, ***P < 0.001 vs PMH, and ^#^P < 0.05 vs OMH.

**Figure 3.** Source-dependent PF4–heparin complexes formation under equipotent and molarity-normalized conditions. (A) SEC analysis of PF4–heparin complexes formation at equipotent anticoagulant concentrations (0.5 IU/mL and 1 IU/mL). (B) Comparison of heparin molar concentrations corresponding to equipotent anticoagulant dosing among PMH, OMH, and BLH. (C) SEC analysis of PF4–heparin complexes formation at an identical heparin molar concentration (3 μM). (D) Competitive ELISA results evaluating HIT antibody recognition by PF4–heparin complexes formed under the identical molar concentration condition (3 μM), expressed as absolute OD values (left) and relative antibody binding percentages (right). Data are presented as mean ± SD (n = 3). Statistical analysis was performed using one-way ANOVA, \**p* < 0.05 vs PMH, ** *p* < 0.01 vs PMH, *** *p* < 0.001 vs PMH; ^##^ *p* < 0.01 vs OMH, ^###^ *p* < 0.001 vs OMH; ns, not significant vs PMH.

**Figure 4.** Source-dependent PF4–LMWH complex formation under equipotent conditions. (A) Comparison of heparin molar concentrations corresponding to equipotent anticoagulant dosing among PME, OME, and BLE. (B) SEC analysis of PF4–LMWH complex formation at equipotent anticoagulant concentrations (0.5 IU/mL and 1 IU/mL). (C) Representative flow cytometry scatter plots (top) displaying the expression of activation markers (CD62P and CD63) following treatment with 1 IU/mL LMWH, with the right quadrants (Q1-LR) designating activated platelet populations. The accompanying bar graph (bottom) presents the quantitative analysis of platelet activation rates (%). Data are presented as mean ± SD (n = 3). Statistical analysis was performed using one-way ANOVA, \**p* < 0.05 vs PME, ** *p* < 0.01 vs PME, *** *p* < 0.001 vs PME; ^##^ *p* < 0.01 vs OME, ^###^ *p* < 0.001 vs OME; ns, not significant vs PME.

**Figure 5.** Differential PF4 Binding of LMWH Oligosaccharides via PF4 Affinity Chromatography. (A) Total ion chromatograms (TICs) of intact oligosaccharide compositions of PME, OME, and BLE analyzed by HILIC–MS. (B) Total ion chromatograms (TICs) from HILIC–MS analysis of PF4 affinity chromatography fractions eluted at physiological ionic strength (0.15 M NaCl) for PME, OME, and BLE. (C) Total ion chromatograms (TICs) from HILIC–MS analysis of PF4 affinity chromatography fractions eluted at near-physiological ionic strength (0.4 M NaCl) for PME, OME, and BLE. (D) Relative quantification of dp6 oligosaccharides in the PF4 affinity chromatography eluates obtained at 0.15 M and 0.4 M NaCl. (E) Relative quantification of dp8 oligosaccharides in the PF4 affinity chromatography eluates obtained at 0.15 M and 0.4 M NaCl. (F) Relative abundance of N-acetylated (1Ac) species among dp6 and dp8 oligosaccharides from different LMWH sources. (G) Quantitative comparison of 1Ac-containing versus non-acetylated oligosaccharides in the flow-through fractions collected at 0.15 M NaCl.

**Figure 6.** Molecular docking analysis reveals a critical role of N-sulfation in PF4–LMWH interactions. (A) The interaction between fully sulfated dp6 and PF4 tetramer. (B) The interaction between reducing end 1Ac dp6 and PF4 tetramer.

## Notes

### Competing Interest Statement

The authors have declared no competing interest.

